# Conserved bacterial genomes from two geographically distinct peritidal stromatolite formations shed light on potential functional guilds

**DOI:** 10.1101/818625

**Authors:** Samantha C. Waterworth, Eric W. Isemonger, Evan R. Rees, Rosemary A. Dorrington, Jason C. Kwan

## Abstract

Stromatolites are complex microbial mats that form lithified layers and ancient forms are the oldest evidence of life on earth, dating back over 3.4 billion years. Modern stromatolites are relatively rare but may provide clues about the function and evolution of their ancient counterparts. In this study, we focus on peritidal stromatolites occurring at Cape Recife and Schoenmakerskop on the southeastern South African coastline. Using assembled shotgun metagenomic data we obtained 183 genomic bins, of which the most dominant taxa were from the Cyanobacteriia class (Cyanobacteria phylum), with lower but notable abundances of bacteria classified as Alphaproteobacteria, Gammaproteobacteria and Bacteroidia. We identified functional gene sets in bacterial species conserved across two geographically distinct stromatolite formations, which may promote carbonate precipitation through the reduction of nitrogenous compounds and possible production of calcium ions. We propose that an abundance of extracellular alkaline phosphatases may lead to the formation of phosphatic deposits within these stromatolites. We conclude that the cumulative effect of several conserved bacterial species drives accretion in these two stromatolite formations.

**ORIGINALITY-SIGNIFICANCE:** Peritidal stromatolites are unique among stromatolite formations as they grow at the dynamic interface of calcium carbonate-rich groundwater and coastal marine waters. The peritidal space forms a relatively unstable environment and the factors that influence the growth of these peritidal structures is not well understood. To our knowledge, this is the first comparative study that assesses species conservation within the microbial communities of two geographically distinct peritidal stromatolite formations. We assessed the potential functional roles of these communities using genomic bins clustered from metagenomic sequencing data. We identified several conserved bacterial species across the two sites and hypothesize that their genetic functional potential may be important in the formation of pertidal stromatolites. We contrasted these findings against a well-studied site in Shark Bay, Australia and show that, unlike these hypersaline formations, archaea do not play a major role in peritidal stromatolite formation. Furthermore, bacterial nitrogen and phosphate metabolisms of conserved species may be driving factors behind lithification in peritidal stromatolites.

## INTRODUCTION

Stromatolites are organo-sedimentary structures that date back more than 3.4 billion years, forming the oldest fossils of living organisms on Earth (Dupraz *et al*., 2009). A more recent discovery of stromatolite-like structures in Greenland suggests that the structures may date as far back as 3.7 - 3.8 billion years (Nutman *et al*., 2016), however, their biogenic origin remains under debate (Witze, 2016). The emergence of Cyanobacteria in stromatolites approximately 2.3 billion years ago initiated the Great Oxygenation Event that fundamentally altered the Earth’s redox potential and resulted in an explosion of oxygen-based and multicellular biological diversity (Soo *et al*., 2017). Ancient stromatolites could provide insight into how microorganisms shaped early eukaryotic evolution. Unfortunately ancient microbial mats are not sufficiently preserved for identification of these microbes and individual bacteria cannot be classified more specifically than phylum Cyanobacteria due to morphological conservatism (Awramik, 1992; Dupraz *et al*., 2009). The study of extant stromatolite analogs may therefore help to elucidate the biological mechanisms that led to the formation and evolution of their ancient ancestors. Modern stromatolites are formed through a complex combination of both biotic and abiotic processes. The core process revolves around the carbon cycle where bacteria transform inorganic carbon into bioavailable organic carbon for respiration. Bacterial respiration in turn results in the release of inorganic carbon, which, under alkaline conditions, will bind cations and precipitate primarily as calcium carbonate (Dupraz *et al*., 2009). This carbonate precipitate, along with sediment grains, can then become trapped within bacterial biofilms forming the characteristic lithified layers.

Alteration of the pH and subsequently, the solubility index (SI), may promote mineralization or dissolution of carbonate minerals through microbial cycling of redox sensitive compounds such as phosphate, nitrogen, sulfur and other nutrients within the biofilm. This in turn regulates the rate of carbonate accretion and stromatolite growth. Particularly, photosynthesis and sulfate reduction have been demonstrated to increase alkalinity thereby promoting carbonate accretion, resulting in the gradual formation of lithified mineral layers (Dupraz *et al*., 2009). In some stromatolite formations such as those of Shark Bay, Australia, there is abundant genetic potential for both dissimilatory oxidation of sulfur (which may promote dissolution under oxic conditions and precipitation under anoxic conditions) and dissimilatory reduction of sulfate (which promotes precipitation) (Gallagher *et al*., 2012; Casaburi *et al*., 2016; Wong *et al*., 2018).

The biogenicity of stromatolites has been studied extensively in the hypersaline and marine formations of Shark Bay, Australia and Exuma Cay, Bahamas, respectively (Khodadad and Foster, 2012; Babilonia *et al*., 2018). The presence of Archaea has been noted in several microbial mat and stromatolite systems (Casaburi *et al*., 2016; Balci *et al*., 2018; Medina-Chávez *et al*., 2019), particularly in the stromatolites of Shark Bay, where they are hypothesized to potentially fulfill the role of nitrifiers and hydrogenotrophic methanogens (Wong *et al*., 2017). Although Cyanobacteria, Proteobacteria and Bacteroidetes appear to be abundant in both marine and hypersaline systems, Cyanobacteria are proposed to be particularly vital to these formations through the combined effect of biofilm formation, carbon fixation, nitrogen fixation and endolithic (boring) activity (Macintyre *et al*., 2000; Khodadad and Foster, 2012; Casaburi *et al*., 2016; Babilonia *et al*., 2018).

Peritidal tufa stromatolite systems are found along the southeastern coastline of South Africa (SA). They are geographically isolated, occurring at coastal dune seeps separated by stretches of coastline (Smith *et al*., 2018). In these systems, stromatolite formations extend from freshwater to intertidal zones and are dominated by Cyanobacteria, Bacteroidetes and Proteobacteria (Perissinotto *et al*., 2014). The stromatolites are impacted by fluctuating environmental pressures caused by periodic inundation by seawater, which affects the nutrient concentrations, temperature and chemistry of the system (Rishworth *et al*., 2016). These formations are characterized by their proximity to the ocean, where stromatolites in the upper formations receive freshwater from the inflow seeps, middle formation stromatolites withstand a mix of freshwater seepage and marine over-topping, and lower formations are in closest contact with the ocean (Perissinotto *et al*., 2014). The stromatolite formations at Cape Recife and Schoenmakerskop are exposed to both fresh and marine water that has little dissolved inorganic phosphate and decreasing levels of dissolved inorganic nitrogen (Cape Recife: 82 - 9 µM, Schoenmakerskop: 424 - 14 µM) moving from freshwater to marine influenced formations (Rishworth, Perissinotto, Bird, *et al*., 2017). Microbial communities within these levels therefore likely experience distinct environmental pressures, including fluctuations in salinity and dissolved oxygen (Rishworth *et al*., 2016). While carbon predominantly enters these systems through cyanobacterial carbon fixation, it is unclear how other members of the stromatolite-associated bacterial consortia influence mineral stratification resulting from the cycling of essential nutrients such as nitrogen, phosphorus and sulfur. Since peritidal stromatolites exist in constant nutritional and chemical flux with varying influence from the fresh and marine water sources, they present an almost ideal *in situ* testing ground for investigating which microbes are consistently present despite fluctuations in their environment. Identification of conserved bacterial species across both time and space and across varied environmental pressures would suggest that these bacteria are not only robust but likely play important roles within the peritidal stromatolite consortia.

Previous studies of stromatolites collected from Lake Clifton, Pavilion Lake, Clinton Creek, Highborne Cay and Pozas Azules have investigated the overall bacterial composition using 16S rRNA gene analysis and/or overall potential gene function of unbinned metagenomic datasets per bacterial taxa using tools such as MG-RAST (Keegan *et al*., 2016) to gain insight into the potential roles of the different bacterial groups (Mobberley *et al*., 2015; Centeno *et al*., 2016; Gleeson *et al*., 2016; Ruvindy *et al*., 2016; Warden *et al*., 2016; White *et al*., 2016). However, the presence of all genes required for a complete functional pathway within a collection of bacteria does not necessarily mean that the cycle can take place since they may not be present within a single organism. Therefore, in order to assess the potential for functional roles, individual bacterial genomes must be investigated. To date, only two studies have successfully binned individual bacterial genomes from culture-independent, metagenomic data originating from stromatolites from Shark Bay (Australia) (Wong *et al*., 2018) and Socompa Lake (Argentina) (Kurth *et al*., 2017). The latter study obtained four high-quality bins (in accordance with MIMAG standards defined by (Bowers *et al*., 2017)) and analyzed three which were believed to have carried genes from several closely-related genomes. Extracted 16S rRNA sequences were in conflict with whole-genome taxonomic classifications (Kurth *et al*., 2017). The Shark Bay study obtained a total of 550 binned genomes, of which 87 (15.8%) were of medium to high quality (Bowers *et al*., 2017; Wong *et al*., 2018) and the data provided information on a number of potentially important processes that may contribute to the formation and maintenance of the hypersaline stromatolites. However, the study did not identify key microbial architects within these biogenic structures.

Using a metagenomic approach, we have sought to gain insight into the foundational bacteria responsible for metabolic processes that potentially result in formation of pertidal South African stromatolites. We obtained and annotated 183 putative bacterial metagenome-assembled genomes (MAGs), (of which 112 (61%) were of medium to high quality) from samples of two geographically isolated sites near Port Elizabeth, South Africa. We identified several temporally and spatially conserved bacterial species and functional gene sets, that are likely central in establishing and maintaining peritidal stromatolite microbial communities.

## RESULTS AND DISCUSSION

Two geographically isolated peritidal stromatolite sites, Cape Recife and Schoenmakerskop, which are 2.82 km apart, were chosen for this study. These two sites have been extensively characterized with respect to their physical structure, nutrient and chemical environment (Perissinotto *et al*., 2014; Rishworth *et al*., 2016, 2019; Rishworth, Perissinotto, Bornman, *et al*., 2017; Dodd *et al*., 2018). The sites experience regular shifts in salinity due to tidal overtopping and groundwater seepage (Rishworth *et al*., 2019). Comparison to a site with groundwater seepage but no stromatolite growth, has shown that the growth of peritidal stromatolites is promoted within this region by decreased levels of wave action, higher water alkalinity and decreased calcite and aragonite saturation (Dodd *et al*., 2018). Furthermore, stromatolite growth is inhibited by increased levels of salinity, as lithified structures are not observed in both the marine waters and in deeper portions of formation pools, with higher salinity levels (Dodd *et al*., 2018).

Stromatolite formations at both sites begin at a freshwater inflow and end before the subtidal zone (Fig. 1) and are exposed to different levels of tidal disturbance. Samples for this study were collected from the upper stromatolites at Cape Recife and Schoenmakerskop in January and April 2018 for comparisons over time and geographic space. Additional samples were collected in April 2018 from middle and lower formations for extended comparison across the two sites (Fig. 1). For a detailed perspective of depth and the differentiation of the sampled zones a 3-dimensional rendering of the sample sites was constructed by a 3rd party (Caelum Technologies^⍰^) using a combination of drone-based photogrammetry and geographic mapping using differential GPS (Fig. 1). Stromatolite formations were classified according to their tidal proximity as defined in previous studies (Perissinotto *et al*., 2014; Rishworth *et al*., 2016; Rishworth, Perissinotto, Bornman, *et al*., 2017; Dodd *et al*., 2018). Upper formations occur in the supratidal zone where they are constantly exposed to fresh water flowing from dune seeps. These formations will only be exposed to seawater during spring tides or extreme storm surges. The water from upper formations feeds into large pools, which in turn feed into the lower portion of the system. In the upper-middle intertidal zone, the formations form a slope into large pools where they will only receive seawater during peak high tide. Lower formations occur in areas where saline or brackish conditions are predominant. In Schoenmakerskop the lower zone is located in a semi stagnant pool which is frequently overtopped, whilst in the lower zone of Cape Recife, formations create a low flowing slope which ends at the subtidal zone. All sampling was conducted at low tide. Sample abbreviations and MAG prefixes used throughout this study correspond to the site and region from which they were sampled e.g. “SU” identifies the sample as originating from <u>S</u>choenmakerskop, <u>U</u>pper formation in April (Table S1). Samples prefixed with a “C” identify samples collected in January e.g. “CSU” identifies the sample as originating from <u>S</u>choenmakerskop, <u>U</u>pper formation in January (Table S1).

**Figure 1.**
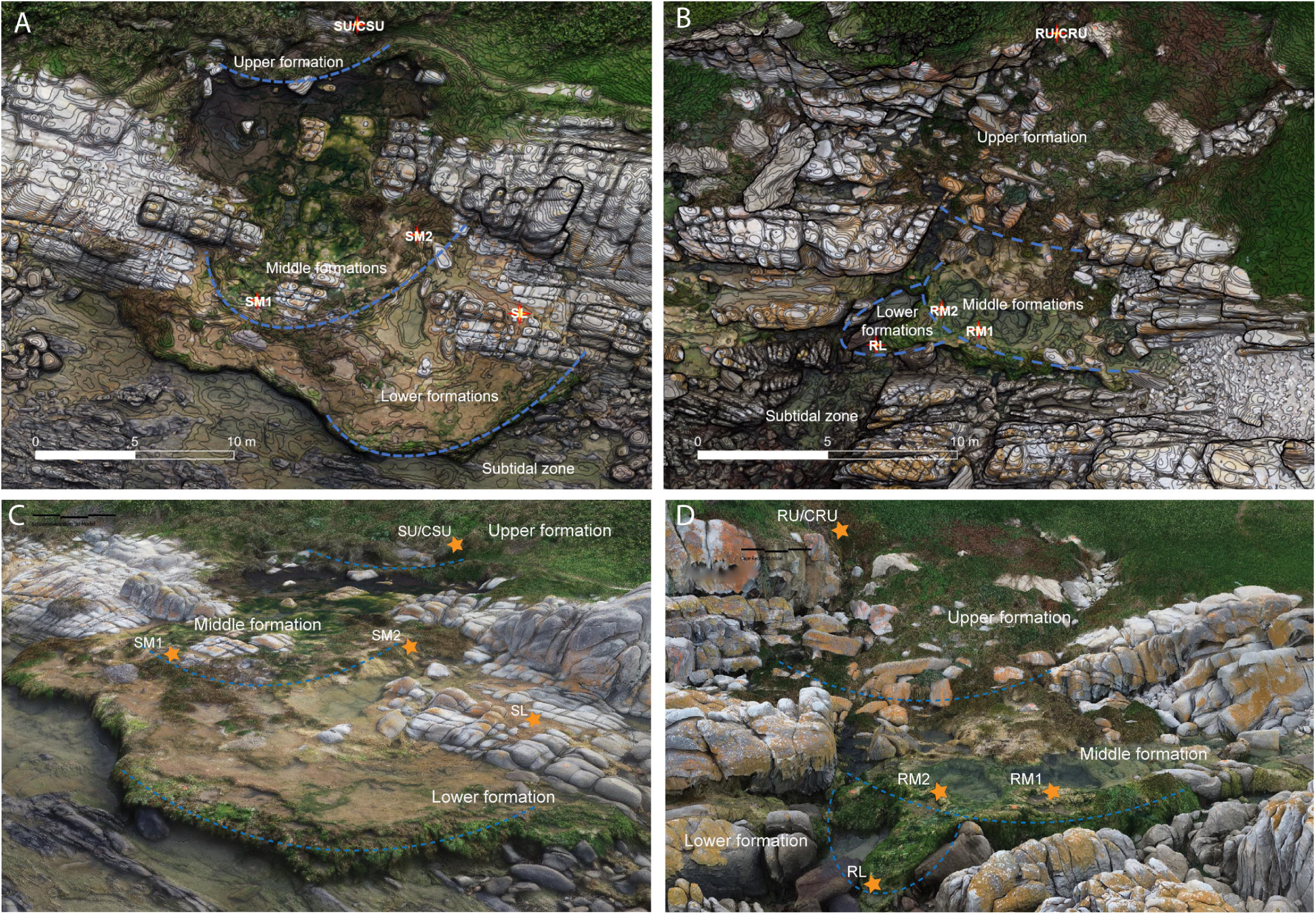
Stromatolites were collected from four different points at two different sites along the SA coastline. (A) Aerial view of sampling locations within the Schoenmakerskop site and (B) the Cape Recife site. Contour lines represent elevation at 5cm increments. Samples were collected from upper, middle and lower formations. The boundaries of which are indicated with a dotted blue line. (C - D) A 3-dimensional rendering of the sample sites at Schoenmakerskop and Cape Recife respectively constructed by a 3rd party (Caelum Technologies^⍰^) using a combination of drone-based photogrammetry and geographic mapping using differential GPS. Samples were collected from upper, middle and lower formations; the boundaries of which are indicated with a dotted blue line. Abbreviations are as follows: CSU: Schoenmakerskop Upper Jan SU: Schoenmakerskop Upper April, CRU: Cape Recife Upper (Jan), RU: Cape Recife Upper (April), SM1: Schoenmakerskop Middle 1 (April), SM2: Schoenmakerskop Middle 2 (April), RM1: Cape Recife Middle 1 (April), RM2: Cape Recife Middle 1 (April), SL: Schoenmakerskop Lower (April), RL: Cape Recife Lower (April).

### Phylogenetic distribution of microbial communities

We assessed the diversity and structure of the bacterial communities in triplicate samples taken from upper, middle and lower formations at Cape Recife and Schoenmakerskop using 16S rRNA gene amplicon sequence analysis. All communities were dominated by Cyanobacteria, Bacteroidetes, Alphaproteobacteria, Gammaproteobacteria and other unclassified bacteria (Fig. 2A), in agreement with a previous study at Schoenmakerskop (Perissinotto *et al*., 2014). Bacterial communities at each sampling site were statistically different from one another (Fig. 2B; ANOSIM: R = 0.976, p = 0.01 permutations = 999), whilst pairwise Kendall rank correlation tests showed that replicates were not statistically different from one another (pairwise p-values < 2.2 × 10^-16^).

**Figure 2.**
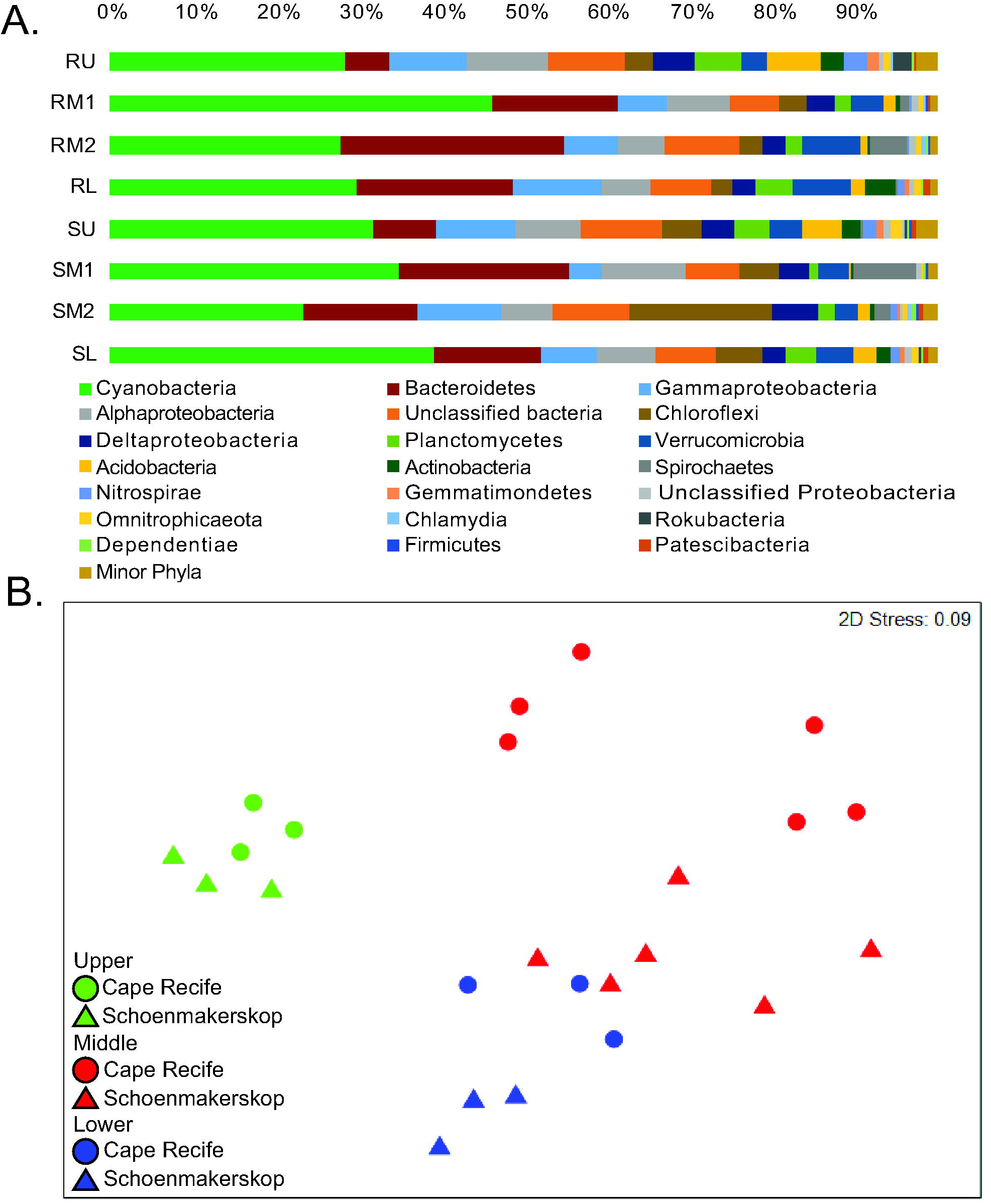
Distribution and abundance of bacterial taxa in stromatolite formations based on 16S rRNA gene fragment amplicon libraries. (A) Phylogenetic classification and average relative abundance (n=3) of dominant phyla in different sample sites indicated that all stromatolite samples are dominated by Cyanobacteria, Bacteroidetes, Alpha- and Gammaproteobacteria. (B) OTU abundance was used to cluster stromatolite biological replicates using Bray-Curtis non-dimensional scaling and showed statistically significant clustering of replicate samples from each of the sampled regions. Samples were isolated from upper (green), middle (red) and lower (blue) stromatolites formations in Cape Recife (triangles) and Schoenmakerskop (circles).

### Trends observed in metagenomes of stromatolite samples from Cape Recife and Schoenmakerskop

To characterize the metabolic potential within stromatolites, we generated shotgun metagenomic libraries from 8 samples representative of the upper, middle and lower formations at both sites in April 2018, as well as an additional two samples from the upper formations of each site, collected 4 months previously in January 2018 (Table S1). Following assembly of raw sequence reads, gene coverages were normalized to the length-weighted average coverage per sample, revealing a high abundance of genes encoding phosphate transport (*pstSCAB*), phosphate uptake regulation (*phoURBP*) and alkaline phosphatases (*phoADX*) were observed across the board (Fig. 3). Additionally, genes involved in phosphonate metabolism (*phnCDEFGHIJKLM)* were abundant in three of the four middle formations but were absent in upper formations. The concentration of soluble phosphorus, although relatively low, is highest in water surrounding the middle formations, as both fresh seep water and ocean overtopping contribute to the total phosphate (Dodd *et al*., 2018). It is thus unsurprising then, that the greatest abundance of phosphate-metabolism genes are found in the middle formations. The high abundance of genes encoding phosphate-metabolizing enzymes, may also be indicative of how stromatolite communities cope with low dissolved inorganic phosphorus in their environment. Genes encoding alkaline phosphatase *phoX*, were the most abundant among the phosphatases observed in these stromatolite samples, and represent a calcium-dependent enzyme that can function at low substrate concentrations on a broad range of C-O-P substrates (Zaheer *et al*., 2009). The phosphonate transporter genes (*phnCDE*) are more prevalent than the genes encoding the C-P lyase (*phnGHIJLM*) required for phosphonate degradations (Fig. 3). These transporters have also been implicated in the transport of inorganic phosphate, in addition to phosphonates (Stasi *et al*., 2019), and the discrepancy between transporters and metabolic genes may suggest an additional role of phosphate uptake in these systems. Overall gene abundances indicated negligible presence of canonical dissimilatory sulfate reduction/oxidation via *aprAB* and *dsrAB* encoded enzymes. Similarly, there were low abundances of genes associated with sulfonate metabolism. There was an abundance of genes associated with assimilatory sulfate reduction, but the low abundance or absence of *cysC*, a gene encoding a key enzyme in this pathway, indicated that this pathway may not be complete. Genes associated with assimilatory nitrate reduction (*narB*, *nirA*) were the most prevalent markers of nitrogen metabolism. All sites also contained a number of genes associated with dissimilatory nitrate reduction where cytoplasmic NADH-dependent nitrate reductase *nirBD* appeared to be favored over periplasmic cytochrome c nitrate reductase *nrfAH*. Interestingly nitrogen fixation genes (*nifDHK*) appear to be more abundant in the middle stromatolite formations of Schoenmakerskop (Fig. 3).

**Figure 3.**
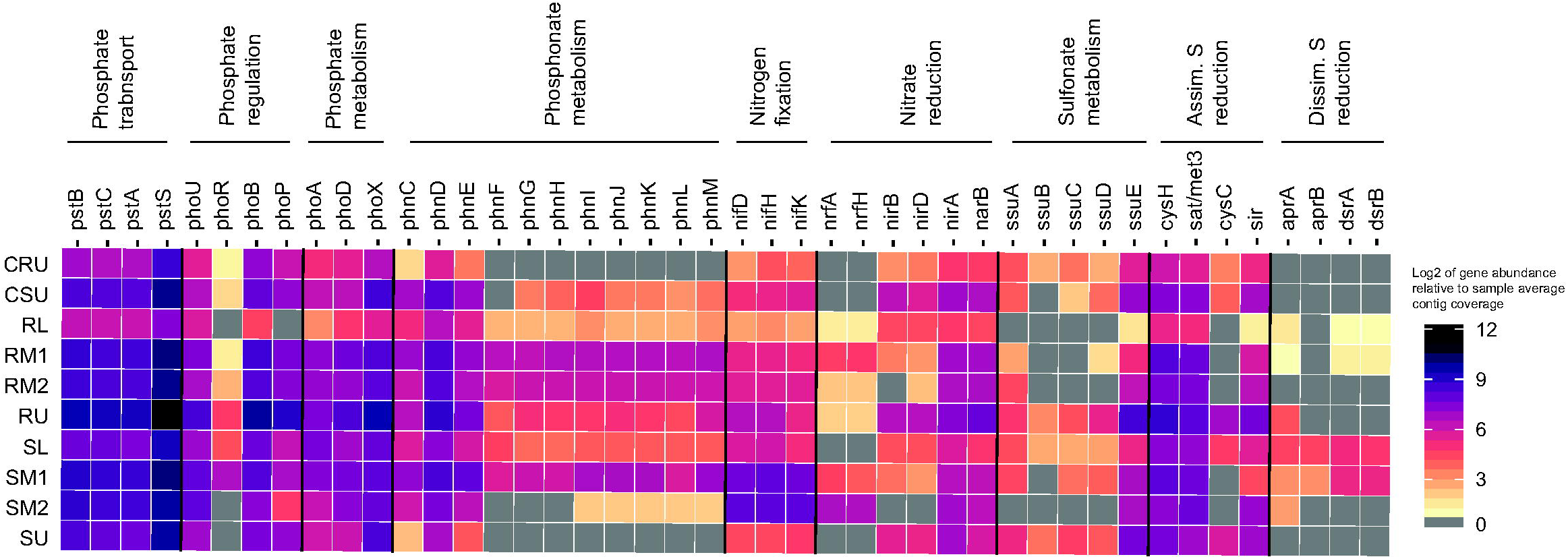
Summary of phosphate, nitrogen and sulfate transport and metabolism genes in overall metagenomic data from stromatolites in Schoenmakerskop and Cape Recife upper, middle and lower formations respectively. Gene abundance is expressed relative to the length-weighted average contig coverage per sample and transformed using log_2_ scaling. Note: On the color scale, grey indicates genes which were not detected in the respective sample. Abbreviations are as follows: CSU: Schoenmakerskop Upper Jan SU: Schoenmakerskop Upper April, CRU: Cape Recife Upper (Jan), RU: Cape Recife Upper (April), SM1: Schoenmakerskop Middle 1 (April), SM2: Schoenmakerskop Middle 2 (April), RM1: Cape Recife Middle 1 (April), RM2: Cape Recife Middle 1 (April), SL: Schoenmakerskop Lower (April), RL: Cape Recife Lower (April).

### Binning and phylogenetic classification of putative genomes

The 10 assembled metagenomes were binned using Autometa (Miller *et al*., 2019), resulting in a total of 183 bacterial genome bins (Table S1). Using relative coverage per sample as a proxy for abundance, we found that genomes classified within the Cyanobacteriia class were consistently dominant in all collection points, while Alphaproteobacteria, Gammaproteobacteria and Bacteroidia were less abundant but notable bacterial classes (Fig. 4A). This distribution appears to be approximately congruent with abundances observed in the 16S rRNA gene amplicon analyses (Fig. 2).

**Figure 4.**
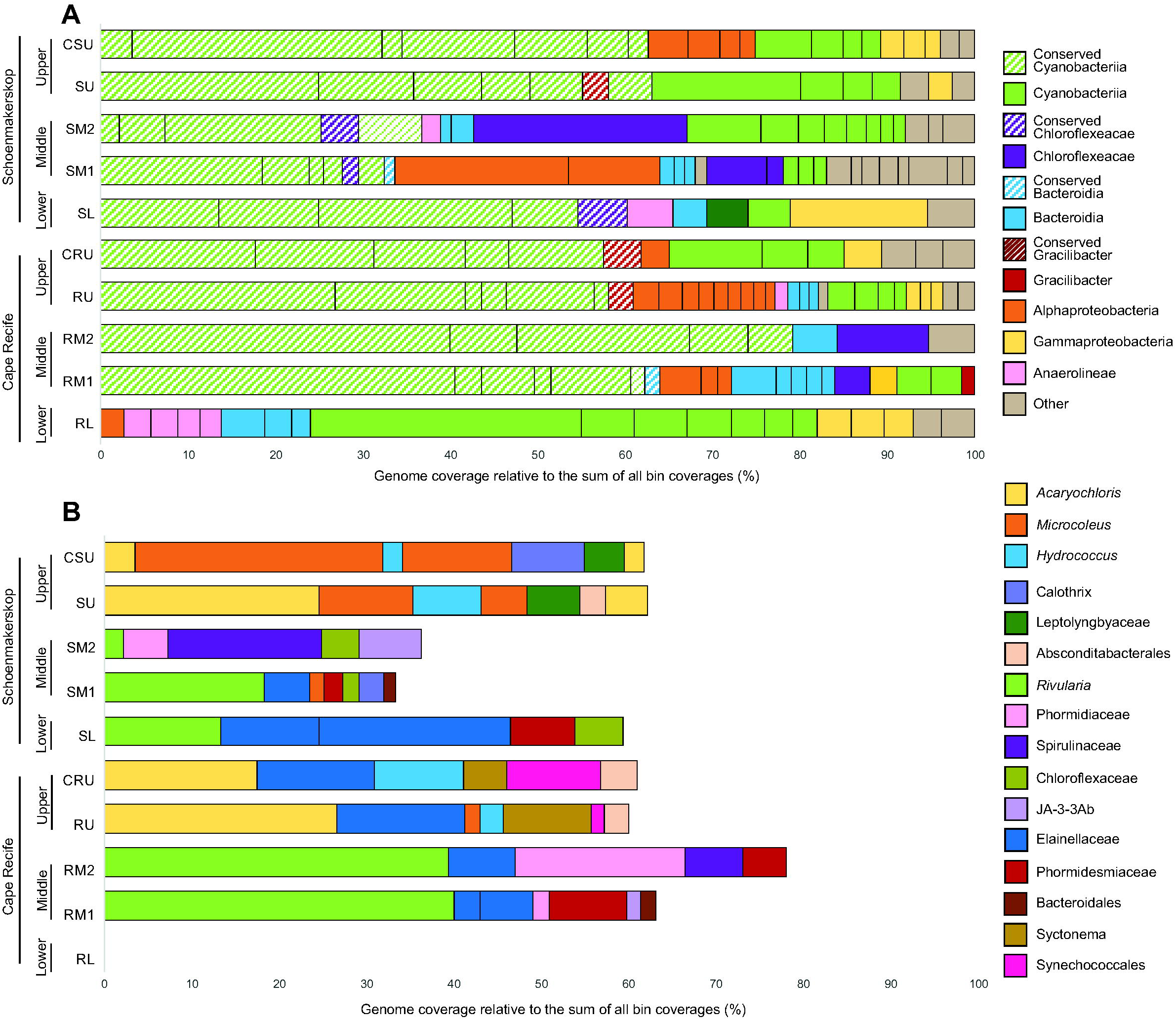
Taxonomic classification of putative genome bins in stromatolites collected from upper/inflow, middle and lower/marine formations of Schoenmakerskop and Cape Recife. Coverage per genome has been used as a proxy for abundance and used to scale the size of individual genome bars, expressed as a percentage of the sum of all genome bin coverages. (A) The coverage of conserved (diagonal lines) and non-conserved (solid colors) bacterial genomes. Taxonomic classification of each genome is indicated by color. (B). Taxonomic classifications of conserved bacterial species in each of the samples, generated with GTDB-Tk (Chaumeil *et al*., 2019).

### Temporal and spatial conservation of bacterial species

We calculated pairwise average nucleotide identity (ANI) between all binned genomes and defined conserved species as genomes sharing more than 97% ANI in two or more of the sampled regions. We identified 16 conserved taxa across the 10 sampled regions, with several species identified in 3 - 5 samples, across Schoenmakerskop and Cape Recife, however no single species was common to all sampled sites (Table S2, Fig. 4A). Conserved species were commonly the most abundant taxa present in each of the samples, accounting for approximately 30 - 80% of the species abundance (Table S2, Fig. 4A). Cyanobacterial species within the *Acaryochloris* and *Hydrococcus* genera and Absconditabacterales family were conserved across upper formations (Table 1, Fig. 4B, Table S3). Seven other species were conserved across the middle formations, including species classified within the *Rivularia* genus and Phormidesmiaceae and Spirulinaceae families (Table 1 and S3, Fig. 4B). Two distinct species within family Elainellaceaee were also conserved: species A was detected only in the upper pools of Cape Recife, whilst species B was conserved across the middle formations of both sites (Table 1 and S3, Fig. 4B).

Also, of interest was the presence of conserved bacterial species (order Absconditabacterales), which are classified under the Patescibacteria phylum. Patescibacteria are unusually small bacteria found in groundwater that produce large surface proteins hypothesized to help them attach to, and exploit the ability of other microorganisms performing nitrogen, sulfur and iron cycling (Herrmann *et al*., 2019). The presence of these conserved bacteria suggests that the inflow water seeps may originate from groundwater.

Conserved *Rivularia*, Elainellaceae, Phormidesmiaceae and Chloroflexaceae species were identified in the lower formation of Schoenmakerskop, but none of the genomes identified in the lower formation of Cape Recife had greater than 97% shared ANI with any other genome bin. The distribution of conserved taxa, wherein the upper and middle formations appear to harbor distinct conserved taxa suggests that the differing nutrient and physical characteristics between upper, middle and lower regions of the stromatolite formation elicit specialization of the conserved bacterial community. The lack of conserved bacterial species across the lower formation of Cape Recife may be due to the choice of sampling site: The lower formation sample taken from Cape Recife is in closer contact with the ocean than the lower formation sample taken from Schoenmakerskop (Fig. 1 C - D). The proximity of the Schoenmakerskop lower formation sample to the ocean was initially thought to be sufficient to delimit it as a lower formation, but following this study it may be reclassified as a middle formation, as the bacterial community composition corresponds more closely with the other four middle formation samples (Fig. 4B).

### Metabolic potential of binned genomes

Oxygenic photosynthesis by cyanobacteria results in rapid fixation of carbon dioxide and an increase in alkalinity (Pace *et al*., 2018). Carbonate ions bind cations such as calcium and are precipitated under alkaline conditions, promoting the growth of stromatolite structures (Dupraz *et al*., 2009). Given their numerical dominance in Cape Recife and Schoenmakerskop stromatolites, and their predicted role in other stromatolites (Dupraz *et al*., 2009), cyanobacteria likely perform carbon sequestration. However, the identity of the bacteria that cycle redox-sensitive sulfur, phosphate, nitrogen and calcium, and subsequently affect the alkalinity and solubility index enabling carbonate precipitation in these stromatolites remains unknown. We inspected PROKKA and KEGG annotations within all stromatolite-associated bacterial genome bins to identify potential metabolic pathways that may promote mineral deposition and accretion (Dupraz *et al*., 2009). The results presented here are summarized in Table 1.

#### Sulfur metabolism

Reduction of sulfate has previously been shown to promote the precipitation of carbonates in the form of micritic crusts in Bahamian and Australian stromatolites (Reid *et al*., 2000; Wong *et al*., 2018) and it has been suggested that microbial cycling of sulfur played an important role in ancient Australian stromatolites, even prior to the emergence of Cyanobacteria (Bontognali *et al*., 2012; Allen, 2016). Amongst the Cape Recife and Schoenmakerskop stromatolite-associated bacteria, the potential capacity for sulfate reduction was confined to only a few genomes (Fig. 5 and Fig. S1). The complete set of genes required for assimilatory sulfate reduction (*sat*/*met3*, *cysC*, *cysH* and *sir* genes) (Santos *et al*., 2015) were recovered in four genomes (Fig. S1), three of which were conserved *Acaryochloris* species (Fig. 5). The abundance of genes associated with assimilatory sulfate reduction in the conserved *Acaryochloris* genome bins accounted for 22 - 100% of gene counts observed in their respective metagenomes (Fig.5). This was calculated as abundance of geneX in binX, as a percentage of geneX abundance in the sample from which the bin was derived (Fig. 5). The greatest abundance genes for uptake and desulfonation of alkanesulfonates (*ssuABCDE*) (Aguilar-Barajas *et al*., 2011; Ellis, 2011) were detected exclusively in conserved *Hydrococcus* species (Fig. 5), which account for 23 - 100% of gene abundance in the respective metagenomes. All genes required for a complete pathway found exclusively in bin RU2_2, but both SU_1_0 and CRU1_1 appear to be missing the *ssuE* gene. The *ssuE* gene is not required for growth using aliphatic sulfonate or methionine substrates, but is required for arylsulfonate metabolism (Kahnert *et al*., 2000). This suggests that these *Hydrococcus* strains are all capable of some form of sulfonate metabolism, but not all can metabolize arylsulfonates (Bin CSU_1_8 is only 37% complete, and therefore of low quality and may be missing gene due to incompleteness). In both cases, these trends are in agreement with the patterns observed in the metabolic potential of the overall metagenome (Fig.3). *Hydrococcus* and *Acaryochloris* species are dominant in upper formations (Fig. 4 and Table S3) and the potential for cumulative removal of hydrogen by sulfate reduction by these species may aid in the creation of an alkaline environment within the system. Seep waters feeding both Cape Recife and Schoenmakerskop have relatively high levels of sulfate (Dodd *et al*., 2018), and it is possible that the *Hydrococcus* and *Acaryochloris* species utilize this nutrient in the upper formations (closest to the seeps) as a selective advantage, simultaneously increasing the alkalinity of the surrounding environment through hydrogen assimilation. There was no evidence for the potential for dissimilatory sulfate reduction in conserved genome bins (Fig. 5). Similarly, amongst all 183 genome bins, only genomes representative of a *Thioiploca* sp. from the Schoenmakerskop lower formation and a *Desulfobacula* sp. from the Schoenmakerskop middle formation included all genes required for dissimilatory sulfate reduction (Fig. S1). The lack of genes associated with conserved or dominant bacterial taxa in the middle and lower pools suggest that either sulfate is not required by the consortia that inhabit the middle/lower formations and is required only by bacteria exposed to ground water, or that insufficient amounts of the sulfate continues into the lower pools where the middle and lower formations are found. This suggests that calcite formation in these peritidal stromatolites may be influenced by processes other than dissimilatory sulfate reduction, in contrast to stromatolite formations in the Cayo Coco lagoonal network, Highborne Cay and Eleuthera Island in the Bahamas (Visscher *et al*., 2000; Dupraz *et al*., 2004; Pace *et al*., 2018).

**Figure 5.**
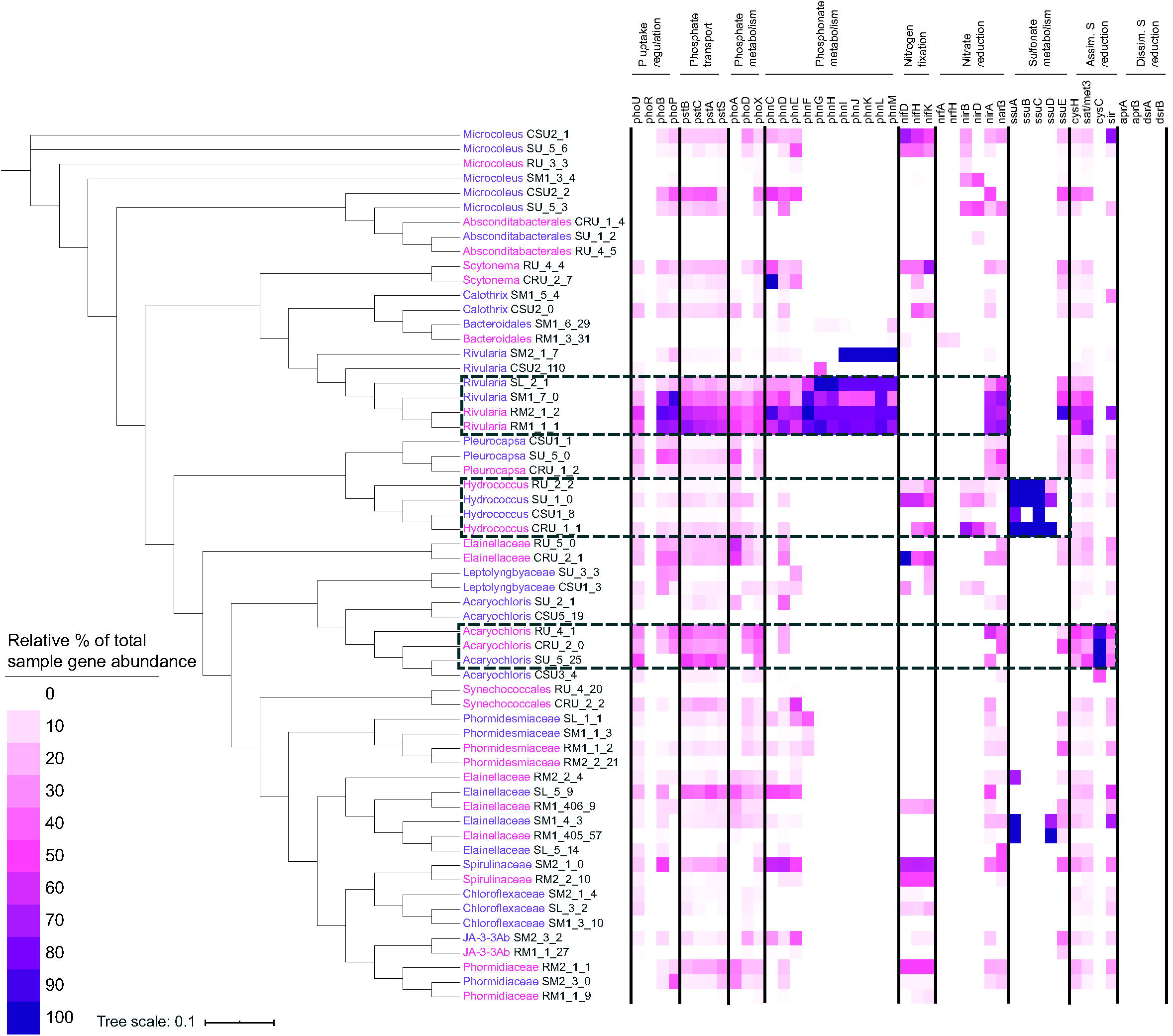
Summary of phosphate, nitrogen and sulfate transport and metabolism genes in conserved genomes from bacteria associated with stromatolites in Schoenmakerskop and Cape Recife upper, middle and lower formations respectively. Relative gene abundance per sample is indicated using a gradient color scale. Relative gene abundance was calculated by dividing coverage-corrected gene abundance in each genome by the coverage-corrected gene abundance from the metagenomic sample from which it came. CSU: Schoenmakerskop Upper Jan SU: Schoenmakerskop Upper April, CRU: Cape Recife Upper (Jan), RU: Cape Recife Upper (April), SM1: Schoenmakerskop Middle 1 (April), SM2: Schoenmakerskop Middle 2 (April), RM1: Cape Recife Middle 1 (April), RM2: Cape Recife Middle 1 (April), SL: Schoenmakerskop Lower (April), RL: Cape Recife Lower (April).

#### Nitrogen metabolism

The reduction of nitrogen, nitrates and nitrites can lead to calcite precipitation (Rodriguez-Navarro *et al*., 2003; Wei *et al*., 2015; Konopacka-Łyskawa *et al*., 2017; Wong *et al*., 2018; Lee and Park, 2019), and the released NH_3_ can react with CO_2_ and H_2_O, to form 2NH^4+^ + CO_3_ (Konopacka-Łyskawa *et al*., 2017). Nitrogen metabolism is proposed to have emerged at approximately the same time as sulfur metabolism, dated to 3.4 billion years ago (Stüeken, 2016), as both processes share similar redox states (Thomazo *et al*., 2011), and ammonium availability during this time may have sustained developing microbial life (Yang *et al*., 2019). Therefore, bacteria that can fix nitrogen or produce ammonia from nitrates/nitrites could potentially promote the growth of stromatolites (Visscher and Stolz, 2005) and may have added to the formation of ancient analogs. Similarly, denitrification will likely lead to mineral dissolution and retardation of stromatolite growth (Visscher and Stolz, 2005). We found that several conserved bacteria carried the genes associated with ferredoxin-dependent assimilatory nitrate reduction (*nirA-narB* genes) (Moreno-Vivián *et al*., 1999), nitrogen fixation (*nifDHK* genes) and dissimilatory nitrite reduction (*nirBD* genes) (Griffith, 2016), all of which result in the production of ammonia (Fig. 5).

The potential for assimilatory nitrate reduction was detected in several genomes, primarily from the middle formations, with particularly high gene abundances of *nirA-narB* genes conserved *Rivularia* sp. (Fig. 5), which are the dominant species in the middle formations (Fig. 4). The potential for dissimilatory reduction of nitrites, via either cytoplasmic *nirBD* or membrane-bound *nrfAH* nitrite reductases, was detected in several genomes across both the upper and middle formations (Fig. S1). It appears that the gene abundances associated with assimilatory nitrate reduction in *Rivularia* sp. account for the majority (29 - 75%, Fig. 5) of *nirA-narB* genes observed in their respective metagenomes (Fig. 3). Among conserved bacteria, the potential for dissimilatory nitrate reduction is relatively low, with *Hydrococcus* sp. carrying the greatest abundance of *nirBD* genes (Fig. 5) which encode cytoplasmic nitrite reductases. Non-conserved *Thioploca, Cyclobacteriaceae* and *Limnothrix* species appear to account for the remainder of the *nirBD* genes observed in their respective metagenomes (Fig. S1). Very few genomes included the *nrfAH* genes which encode membrane-bound nitrite reductases.

Nighttime nitrogen fixation has previously been shown to be a driver of carbonate precipitation in stromatolites (Dupraz *et al*., 2009) and *nif* genes were identified in several conserved bacteria associated with both upper and middle formations: *Hydrococcus* species across both upper formations, *Microcoleus* in the Schoenmakerskop upper formations and Phormidiaceae, Spirulinaceae and Chloroflexaceae species the middle formations (Fig. 5). In addition to the conserved bacteria carrying *nif* genes, non-conserved *Blastochloris* species appeared to account for the majority (10% - 75%) of identified *nifDHK* genes (Fig. S1). The genetic capacity for nitrogen fixation being retained in Schoenmakerskop and Cape Recife was unexpected, given the high concentrations of dissolved inorganic nitrogen (DIN) ranging from 95 - 450 mM at the two sites (Rishworth *et al*., 2016).

### Phosphate and phosphonate metabolism

Phosphatic structures that closely resemble fossilized phosphatic stromatolites have recently been observed within Cape Recife stromatolites (Buttner *et al*., 2019, unpublished). Hydroxyapatite (Ca_5_(PO_4_)_3_(OH)) is more easily precipitated than calcite (CaCO_3_) and the release of PO_4_^3-^ into the biofilm by alkaline phosphatase activity could increase the potential for apatite mineralization, the rate of which may be increased in the presence of an alkaline environment (Gallagher *et al*., 2013). Copies of alkaline phosphatase genes *phoD* and *phoX* are variable within bacterial genomes but the majority of bacteria carry only 1 copy of *phoD* and 1 copy of *phoX* (Ragot *et al*., 2015, 2017). Conserved *Acaryochloris* and *Rivularia* species were particularly notable as they encoded between 2–4 copies of *phoX* (Fig. 5), and carried all genes required for phosphate transport (*pstSCAB*) (Fig. 5). Since the Cape Recife and Schoenmakerskop stromatolites experience limited inorganic phosphate availability (Rishworth *et al*., 2016, 2018; Rishworth, Perissinotto, Bird, *et al*., 2017; Dodd *et al*., 2018), associated bacteria may generate bioavailable phosphate from trapped sediments in biofilms resulting in increased concentrations, similar to cyanobacteria living in phosphate-poor rivers (Wood *et al*., 2015). An increase in both PO_4_^3-^ and Ca^2+^ can result in rapid precipitation of apatite in marine phosphorites, freshwater lakes and in some soil bacteria (Danen-Louwerse *et al*., 1995; Guang-Can *et al*., 2008; Cosmidis *et al*., 2015) and this could account for the phosphatic deposits observed within the SA stromatolites (Buttner *et al*., 2019, unpublished).

Finally, conserved *Rivularia* species in the middle and lower formations and other non-conserved bacterial species harbored the 11 genes required for the transport (*phnCDE*) and lysis (*phnFGHIJKLM*) of phosphonate compounds (Fig. 5) (Metcalf and Wanner, 1993). Phosphonates are characterized by the presence of a carbon-phosphorus bond and are biosynthesized by many organisms (White and Metcalf, 2007). C-P lyase (*phnGHIJLKM*) breaks the C-P bond within phosphonate substrates resulting in a hydrocarbon and inorganic phosphate (White and Metcalf, 2007). The inorganic phosphate may then be used within the bacterial cell or released aiding in accretion through increased ion concentration (Rott *et al*., 2018). Conversely, phosphonates can prevent precipitation of calcite by binding to crystal growth sites (Kan *et al*., 2005) and the degradation of these compounds within stromatolite formations may prevent chemical inhibition of stromatolite growth. Given the low availability of phosphate in these systems, phosphonates may also provide an auxiliary source of bioavailable phosphate. The potential for phosphonate degradation was also observed in 8 genomes from the Shark Bay stromatolites (Wong *et al*., 2018). The conservation of phosphonate metabolism across stromatolites would indicate that these bacterial processes may be important within the generalized stromatolite system.

### Archaeal genomes in Cape Recife and Schoenmakerskop stromatolites

Genome-resolved studies of hypersaline stromatolites in Shark Bay revealed that a large proportion of the microbial community consisted of archaeal species (Wong *et al*., 2017, 2018). Scrutiny of the contigs from Cape Recife and Schoenmakerskop showed that none of the datasets comprised more than 1.5% archaeal genes. Assessment of contigs clustered into the Archaeal kingdom bins from the upper and middle formations showed little evidence for the presence of Archaea. However, two low-quality genomes were obtained from the two lower formations SL_Arch_1 and RL_Arch_2, which were classified as Woesearchaeales (order) and *Nitrosoarchaeum* (genus) respectively (Table S3). Alignment of the 16S rRNA gene obtained from SL_Arch_1 (Table S3) against the nr database showed that it shared the greatest sequence homology with an uncultured archaeon clone GWA2 (87% identity). Similarly, analysis of the 16S rRNA gene sequence from RL_Arch_2 (Table S3) indicated that it shared greatest sequence homology with an uncultured archaeon clone 027 (99.79% identity). The coverage of these genomes is among the lowest in each of the lower formation samples (Table S3) and would suggest a low numerical abundance of these archaea. It is compelling that the only Archaea detected in this study are derived from the formations most influenced by saline waters, considering that the hypersaline stromatolites of Shark Bay comprise several archaeal species (Wong *et al*., 2017, 2018), and it would be intriguing to identify whether salinity has an effect on the Archaeal composition of stromatolite-associated microbiota. However, as there are only two, low quality, non-conserved, relatively low-abundance genomes derived from Archaea, it is likely that this kingdom is less important in the stromatolites found at Cape Recife and Schoenmakerskop.

### Key bacteria for stromatolite formation at Cape Recife and Schoenmakerskop

Several bacterial groups may be key players in the formation of stromatolites in Cape Recife and Schoenmakerskop, with conserved bacterial species potentially acting as major contributors (Table 1). There were distinct functional abilities between the bacterial communities associated between the upper and middle formations. The consortia in the upper formations were dominated by conserved *Hydrococcus* and *Acaryochloris* species, which appeared to be capable of sulfate and sulfonate metabolism, whilst the middle formations were dominated primarily by conserved *Rivularia* species, which carry the genetic capacity to metabolize various forms of phosphate substrates and potentially reduce nitrate. The lack of all genes required for phosphonate metabolism in *Rivularia* species in Schoenmakerskop Middle 2 (SM2) was unexpected but may be explained by the lower abundance of *Rivularia* in this sample, resulting in lower quality of genome (Table S3). Similarly, *Hydrococcus* bin CSU_1_8 was of low quality (Table S3) and as a result appeared to be missing 3 of the genes required.

We propose the redox potential and solubility index in these stromatolites is potentially influenced by conserved bacteria generating sulfide (*Acaryochloris* and *Hydrococcus* species) and ammonia ions (*Rivularia* species) for the rapid precipitation of carbonate compounds and subsequent growth of stromatolite structures. The presence of *Microcoleus* species (Phormidiaceae family) and an unidentified genus of Phormidiaceae in both upper and middle formations is notable, as previous studies have found that lamellar precipitation of carbonate occurs on the filaments of cultured Phormidiaceae bacteria isolated from freshwater tufa structures (Payandi-Rolland *et al*., 2019). We further propose an abundance of alkaline phosphatases in several conserved species, as well as the abundance of genes associated with phosphonate degradation in *Rivularia* species, may result in increased local inorganic phosphate concentrations. This free inorganic phosphate may be incorporated into the phosphatic crusts observed in these stromatolites (Buttner *et al*., 2019, unpublished).

The identification of conserved bacteria performing potentially important roles within stromatolites of both Cape Recife and Schoenmakerskop may provide insight into what cyanobacterial species may have played a key role in the formation of ancient phosphatic stromatolites that formed in shallow marine and peritidal environments (Misi and Kyle, 1994; Drummond *et al*., 2015; Buttner *et al*., 2019, unpublished; Shiraishi *et al*., 2019). However, transcriptomics, nutrient uptake and additional hydrochemistry would be required to determine if the proposed role of the conserved bacteria is valid. Similarly, future studies will be needed to determine whether physical (e.g. flow rate) or chemical factors (e.g. nutrient availability) result in our observed difference in functional potential between the different formations.

## EXPERIMENTAL PROCEDURES

### Sample collection and DNA isolation

Samples were collected from Schoenmakerskop (34°02’28.2″S 25°32’18.6″E) and Cape Recife (34°02’42.1″S 25°34’07.5″E) at low tide, from the water surface level. Samples approximately 1cm deep were collected for 16S rRNA analysis in July 2019, from upper, middle and lower stromatolite formations (Fig. 1) in triplicate, approximately 1 cm apart. Sample cores for metagenomic shotgun sequencing were collected in April 2018, approximately 1cm deep, from upper, middle and lower stromatolite formations. Additional samples were collected in January 2018 from only the upper formations for metagenomic sequencing. All samples were stored in RNAlater and flash-frozen until delivery to the lab at which point, they were stored at -20 °C. DNA was extracted from ∼1g of sample using Zymo quick DNA Fecal/Soil Microbe Miniprep Kit (Zymo Research, Cat No. D6010) according to the manufacturer’s instructions.

### Amplicon sequence analysis

Kapa HiFi Hotstart DNA polymerase (Roche, Cat No. KK2500) was used to generate amplicon libraries of the V4-V5 region of the 16s rRNA ribosomal subunit gene with the primer pair E517F (5’-GTAAGGTTCYTCGCGT-3’) and E969-984 (5’-CAGCAGCCGCGGTAA-3’) (Matcher *et al*., 2011) from triplicate samples using the following cycling parameters: Initial denaturation at 98 °C; 5 cycles (98 °C for 45 seconds, 45 °C for 45 seconds and 72 °C for 1 minute); 18 cycles (98 °C for 45 seconds, 50 °C for 30 seconds and 72 °C for 1 minute); final elongation step at 72 °C for 5 minutes. PCR products were purified using the Bioline Isolate II PCR and Gel kit (Bioline, Cat. No. BIO-52060). Samples were sequenced using the Illumina Miseq platform. Amplicon library datasets were processed and curated using the Mothur software platform (Schloss *et al*., 2009). Sequences shorter than 200 nucleotides or containing ambiguous bases or homopolymeric runs greater than 7 were discarded. Sequences were classified using Naive-Bayesian classifier against the Silva database (v132) and VSEARCH software (Rognes *et al*., 2016) was used to remove chimeras. Reads that were 97% similar were combined into operational taxonomic units (OTUs) using the Opticlust method (Westcott and Schloss, 2017). OTU abundance values were converted to relative values and the dataset was then transformed by square root and statistically analyzed using the Primer-e (V7) software package (Gorley and Clarke, 2015). Reads were not classified against the Greengenes database, as the creators of Mothur warn against this practice, citing poor alignment quality in the variable regions.

### Metagenomic binning

As 16S rRNA sequence analysis indicated that there was no statistical difference between triplicate samples, and that sample regions are therefore homogenous, a single sample from sites CRU, RU, RM1, RM2, RL, CSU, SU, SM1,

### SM2 and SL (Fig.1) collected in January and April 2018 was used to prepare shotgun DNA libraries that were sequenced using IonTorrent Ion P1.1.17

Chip technology (Central Analytical Facilities, Stellenbosch University, South Africa). Adapters were trimmed and trailing bases (15 nts) were iteratively removed if the average quality score for the last 30 nts was lower than 16, which resulted in approximately 30–45 million reads per sample. Resultant metagenomic datasets were assembled into contiguous sequences (contigs) with SPAdes version 3.12.0 (Bankevich *et al*., 2012) using the -- iontorrent and --only-assembler options with kmer values of 21,33,55,77,99,127. The -- iontorrent option enables a read error correction step performed by IonHammer, correcting homopolymeric runs inherent to IonTorrent sequencing chemistry (Ershov *et al*., 2019).

### Quantification of metabolism-associated genes

All raw data, count tables and scripts used to process the data can be accessed here: https://github.com/samche42/Conserved_Stromatolite_bacteria_manuscript.git Contigs in each of the 10 samples were clustered into kingdom bins using Autometa (Miller *et al*., 2019). Genes on all contigs within these bins were identified using Prodigal v.2.6.3 (Hyatt *et al*., 2010). Genes were then annotated against the KEGG database using kofamscan (Aramaki *et al*., 2019) with output in mapper format. Coverage corrected KO annotation counts were collected using kegg_parser.py found in the GitHub repo listed above. Briefly, a dictionary of collection sites was made with all KO numbers supplied in a KEGG annotation query list (i.e. functional group). Kofamscan output per collection site was parsed and each time an entry matching that within the query list was found, a count for that annotation was increased by the coverage of the contig on which it was located, resulting in a coverage-corrected count table of functional group KO annotations per collection site. In order to assess differences between collection sites, these values were transformed relative to average coverage per sample (weighted by contig length) using weighted_contig_coverage_calculator.py (see GitHub repo) according to Eq.1, where *i* is contig length and *j* is contig coverage. The abundances were then transformed using log2 (Fig. 3).

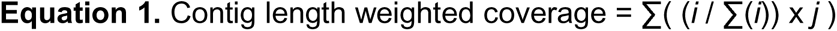

Heatmaps for visualization of the data were created in R using the tidyr, ggplot2 and viridis packages. Scripts can be found in the GitHub repo.

### Clustering of metagenomic data in genome bins

Contigs were then clustered in putative genomic bins using Autometa (https://bitbucket.org/jason_c_kwan/autometa/src/master/, Master branch, commit: a344c28) (Miller *et al*., 2019). Bins were manually curated and resulted in 183 genomic bins. CheckM (Parks *et al*., 2015) was used to assess bin purity and completion using default settings (Table S3). Approximately 37 genomes were of high quality, 75 were of medium quality and 71 were of low quality, in accordance with MIMAG standards defined by (Bowers *et al*., 2017) (Table S3).

### Identificatio**n of conserved taxa**

Conservation of bacterial taxa was calculated using average nucleotide identities (ANIs) of all genomic bins, which were calculated in a pairwise manner using FastANI (Jain *et al*., 2018). All genomic pairs sharing > 97% ANI were subset and considered conserved taxa. Percentage of mapped regions used to calculate ANI is reported in Table S2. Percentage of mapped regions is calculated as the relative number of bidirectional fragments mapping to the total number of query fragments.

### Genome taxonomic classification

Genomes were classified using the standalone GTDB-Tk tool (version 0.3.2) using the classify workflow and Genome Taxonomy Database version 89 (Parks *et al*., 2019). The tool is unable to classify genomes less than 10% complete, as indicated in Table S3. The GTDB-Tk tool uses the Genome Taxonomy Database as a reference for classification, which is based on phylogeny inferred from concatenated protein alignments. This approach enabled the removal of polyphyletic groups and assignment of taxonomy from evolutionary divergence. The resulting taxonomy incorporates substantial changes in comparison to the NCBI taxonomy (Parks *et al*., 2018). Equivalent NCBI taxonomic classifications have been provided for clarity in Table S3.

### Genome taxonomic clustering

Phylogeny of bacterial genomes was inferred using JolyTree (Criscuolo, 2019) with a sketch size of 10 000 (Fig. 5 and Fig. S1). JolyTree infers phylogeny through computation of dissimilarity of kmer sketches, which is then transformed for the estimation of substitution events of the genomes’ evolution (Criscuolo, 2019).

### Genome annotation

Manually curated putative genomic bins were annotated using Prokka version 1.13 (Seemann, 2014), with GenBank compliance enabled. Protein-coding amino-acid sequences from genomic bins were annotated against the KEGG database using kofamscan (Aramaki *et al*., 2019) with output in mapper format. KEGG orthologs were counted and processed as performed quantification of metabolism-associated genes using kegg_parser.py. Relative percentage of genes in individual genomes was calculated by dividing the contig-coverage corrected gene abundance per gene by the total contig-coverage corrected gene abundance for the sample from which it was binned (Fig. 5 and Fig. S1). E.g. The contig-coverage corrected gene abundance of *phoU* in genome bin CRU1_1 was 3.52, and the total contig-coverage corrected gene abundance of *phoU* in the CRU metagenome sample was 29.90253 (See dataset Calculating_rel_perc.xlsx in GitHub repo). Therefore, the relative abundance of *phoU* in genome bin CRU1_1 was: 3.52/29.90 X 100 = 11.79% of total sample gene abundance.

### Archaeal genome binning and identification

Contigs are classified within kingdom bins during Autometa binning. All contigs classified within the Archaea kingdom were given putative taxonomic assignments based on single-copy markers and clustered into bins. No bins could be acquired from the upper and middle formations from either collection site due to few or no contigs being of archaeal origin. Two genomes were obtained from the lower formation of each site respectively (Table S3). Genome quality was assessed using CheckM as described for bacterial bins. 16S rRNA and 23S rRNA sequences were extracted from each genome using barrnap 0.9 (https://github.com/tseemann/barrnap) using the archaeal databases (23S: SILVA-LSU-Arc, 16S: RF01959) as reference. These sequences were then aligned against the nr database using BLASTn (Johnson *et al*., 2008) for putative identification.

## Data availability

Raw 16S rRNA gene amplicon sequence files were uploaded to the NCBI sequence read archive (SRA) database in BioProject PRJNA574289. Raw reads and binned genomes will be deposited in GenBank and respective accession numbers will be included in the accepted version of this manuscript.

## ACKNOWLEDGEMENTS

The authors acknowledge Caro Damarjanan who conducted a pilot study that informed this work. We also wish to thank Karthik Anantharaman (University of Wisconsin-Madison) for his helpful critique.

The authors acknowledge funding from the Gordon and Betty Moore Foundation (Grant number 6920) (awarded to R.A.D and J.C.K.), and grants awarded to R.A.D by the South African National Research Foundation (UID: 87583 and 109680). Development of Autometa in J.C.K.’s laboratory and contributions by E.R.R. were supported by the U.S. National Science Foundation (DBI-1845890). This research was performed in part using the computer resources and assistance of the UW-Madison Center for High Throughput Computing (CHTC) in the Department of Computer Sciences. The CHTC is supported by UW-Madison, the Advanced Computing Initiative, the Wisconsin Alumni Research Foundation, Wisconsin Institutes for Discovery, and the National Science Foundation and is an active member of the Open Science Grid, which is supported by the National Science Foundation and the U.S. Department of Energy’s Office of Science. The authors also acknowledge the Center for High Performance Computing (CHPC, South Africa) for providing computing facilities for bioinformatics data analysis.

The authors declare no conflict of interests.

## TABLE AND FIGURE LEGENDS

Table 1. Conserved bacterial species (ANI > 97%) and their functional potential across upper, middle and lower stromatolite formations at Cape Recife and Schoenmakerskop.

Figure S1. Relative abundance of phosphate, nitrogen and sulfate transport and metabolism genes per genome relative to total gene abundance per sample from Schoenmakerskop and Cape Recife upper, middle and lower formations. Relative gene abundance per sample is indicated using a gradient color scale. Relative gene abundance was calculated by dividing coverage-corrected gene abundance in each genome by the coverage-corrected gene abundance from the metagenomic sample from which it came. CSU: Schoenmakerskop Upper Jan SU: Schoenmakerskop Upper April, CRU: Cape Recife Upper (Jan), RU: Cape Recife Upper (April), SM1: Schoenmakerskop Middle 1 (April), SM2: Schoenmakerskop Middle 2 (April), RM1: Cape Recife Middle 1 (April), RM2: Cape Recife Middle 1 (April), SL: Schoenmakerskop Lower (April), RL: Cape Recife Lower (April).

Table S1. Summary of genomes binned per collected stromatolite sample

Table S2. Conserved bacterial species defined by shared ANI greater than 97% in stromatolite formations from Cape Recife and Schoenmakerskop.

Table S3. Summary of characteristics and taxonomic classifications of genomes binned from shotgun metagenomic data from sampled upper, middle and lower stromatolite formations from Cape Recife and Schoenmakerskop. Bacterial bins are listed first, with the two archaeal bins listed at the bottom.

