## Supplementary figures and images for "Conserved bacterial genomes from two geographically distinct peritidal stromatolite formations shed light on potential functional guilds"

### Figure S1

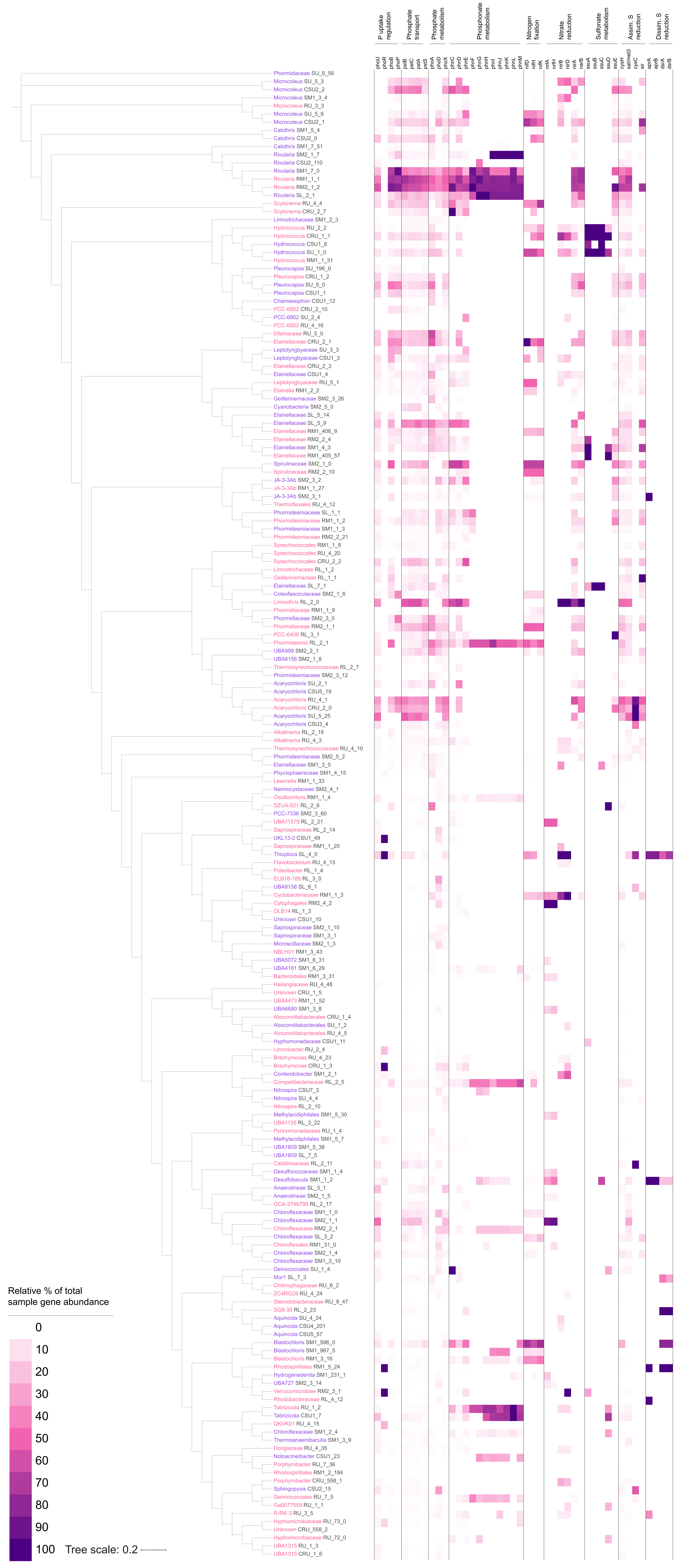
